# Targeting IGF2BP1 Creates a Mitotic Vulnerability in Colorectal Carcinoma

**DOI:** 10.64898/2026.10.22.761131

**Authors:** Bruno Griesler, Sven Hagemann, Robert Winkler, Alice Wedler-Geyer, Claudia Misiak, Danny Misiak, Alexander Rausch, Jana Lützkendorf, Thomas Mueller, Khursheed Ul Islam Mir, Christian Ihling, Florian Wolfgang Otto, Andrea Sinz, Jan-Henning Klusmann, Michael Heuser, Lutz P. Müller, Nadine Bley, Stefan Hüttelmaier

## Abstract

**Background:** Mitotic dysregulation is a common feature of colorectal cancer (CRC), yet the contribution of post-transcriptional mechanisms remain incompletely understood. We investigated how the oncofetal RNA-binding protein IGF2BP1 supports mitotic integrity and exposes therapeutic vulnerabilities in CRC.

**Methods:** We integrated patient cohort analyses with genetic and pharmacological perturbation of IGF2BP1, transcriptomics, proteomics, RNA-binding assays and functional studies. Xenografts assessed tumor formation, while BTYNB–paclitaxel responses were examined in CRC cell lines and patient-derived organoids.

**Results:** IGF2BP1 expression was elevated in CRC, increased with tumor stage, and associated with poor survival and proliferative gene signatures. Genetic depletion or pharmacological inhibition of IGF2BP1 impaired cell viability, induced mitotic abnormalities, and promoted apoptosis, whereas its re-expression enhanced tumor formation *in vivo*. Integrated molecular analyses identified the chromosomal passenger complex (CPC) as a previously unrecognized downstream target of IGF2BP1. IGF2BP1 bound transcripts encoding CPC components and its depletion consistently reduced their expression across multiple cancer cell models. BTYNB disrupted IGF2BP1 association with INCENP, CDCA8, and BIRC5 transcripts and accelerated INCENP mRNA decay. Functionally, INCENP depletion recapitulated the mitotic defects and impaired viability observed upon IGF2BP1 inhibition. Exploiting this mitotic vulnerability, BTYNB synergized with paclitaxel in CRC cell lines and across four independent IGF2BP1-positive patient-derived organoid models.

**Conclusions:** Our findings uncover an IGF2BP1-dependent post-transcriptional regulatory mechanism sustaining CPC expression and mitotic integrity in CRC. Targeting this mechanism exposes a therapeutically exploitable mitotic vulnerability and provides a rationale for combining IGF2BP1 inhibition with microtubule-targeting agents.

## Background

Colorectal cancer (CRC) is one of the most frequently diagnosed and deadliest malignancies worldwide, with approximately 1.9 million new cases and 0.9 million deaths estimated for 2022 (1, 2). Patients with localized disease generally have favorable outcomes following surgical resection, with adjuvant chemotherapy administered according to disease stage and risk factors. However, a substantial proportion of patients are diagnosed with locally advanced or metastatic disease. Treatment options remain limited and include cytotoxic regimens such as FOLFOX and FOLFIRI, anti-angiogenic agents, anti-EGFR antibodies, and immune checkpoint inhibitors for molecularly selected patients (3). Five-year relative survival for metastatic disease remains approximately 13–17% (4–6), which underscores the need for new therapeutic targets based on a mechanistic understanding of CRC biology. CRC is molecularly heterogeneous and can be classified into four consensus molecular subtypes (CMS): CMS1 (MSI immune), CMS2 (canonical), CMS3 (metabolic), and CMS4 (mesenchymal) (7). CMS4 exhibits mesenchymal features, TGF-β activation, and stromal infiltration, whereas CMS2 is characterized by epithelial differentiation, prominent WNT/MYC signaling, pronounced proliferative signatures, and frequent chromosomal instability. Accurate chromosome segregation and cytokinesis depend on the chromosomal passenger complex (CPC), comprising AURKB, INCENP, CDCA8, and BIRC5. Aberrant expression of CPC components has been reported in CRC and associated with tumor progression, but the mechanisms coordinating their expression remain incompletely understood (8, 9).

RNA-binding proteins (RBPs) are central post-transcriptional regulators of oncogenic gene expression, controlling mRNA stability, localization and translation (10). An individual RBP can coordinate networks of functionally related transcripts and thereby amplify oncogenic signals (11). The insulin-like growth factor 2 mRNA-binding protein 1 (IGF2BP1) is a prototypical oncofetal RBP. Its expression is largely restricted to embryonic tissues but is reactivated in many solid tumors, including CRC (12). By binding its target mRNAs, IGF2BP1 stabilizes and enhances the translation of oncogenic transcripts such as MYC/MYCN, YAP1 and E2F1, and thereby promotes proliferation, survival and stemness (13–16). It also contributes to immune evasion, for example by stabilizing CD274/PD-L1 mRNA and indirectly suppressing MHC-I expression (17).

In CRC, IGF2BP1 expression is elevated relative to normal mucosa and increases with pathological stage. High expression is associated with reduced overall survival (18). IGF2BP1 has been reported to sustain Wnt/β-catenin-driven transcriptional programs, stabilize multidrug-resistance transcripts and promote chemotherapy resistance. This highlights its potential as a prognostic biomarker and therapeutic target (12, 19–21). Even so, the full spectrum of IGF2BP1-regulated gene networks in CRC, their impact on tumor progression, as well as the therapeutic value of IGF2BP1 remain incompletely characterized.

BTYNB was identified as the first small-molecule inhibitor of IGF2BP1–RNA binding. It disrupts the interaction of IGF2BP1 with target mRNAs and reduces expression of oncogenic drivers such as MYC/MYCN and E2F1 (15, 16, 22). BTYNB has shown anti-tumor activity in several cancer entities, including ovarian carcinoma, neuroblastoma and leukemia, and has been combined synergistically with other targeted agents (15–17, 23). Enhanced antitumor activity has also been reported in combination with the CDK4/6 inhibitor palbociclib or anti-PD-1 antibodies in ovarian cancer, and with the BET inhibitor mivebresib in MYCN-amplified neuroblastoma (15–17). How BTYNB affects mitotic regulation in CRC, and whether IGF2BP1 expression marks a combination vulnerability in patient-derived models, remains unresolved.

Here, we identify the chromosomal passenger complex (CPC), a key regulator of mitotic progression and chromosome segregation, as a previously unrecognized downstream target of IGF2BP1 in CRC. Using genetic and pharmacological approaches, we demonstrate that IGF2BP1 sustains CPC expression and mitotic integrity, while its depletion or inhibition impairs tumor cell viability. Importantly, pharmacological inhibition of IGF2BP1 synergizes with paclitaxel in CRC cell lines and across four independent IGF2BP1-positive patient-derived organoid models, revealing a therapeutically exploitable mitotic vulnerability.

## Methods

### Analysis of public patient data

Bulk RNA-seq data (TPM) from TCGA-COAD and three CRC cohorts (GSE220148, GSE107422, E-MTAB-12862 [Nunes et al.]) were downloaded from the R2: Genomics Analysis and Visualization Platform (https://r2platform.com). Only histologically confirmed adenocarcinomas were included; mucinous adenocarcinomas and samples without histological subtype were excluded (cohort sizes in Fig. 1D). IGF2BP1 expression was compared between tumor (n = 397) and non-tumor tissue (n = 39) and between stages I/II (n = 222) and III/IV (n = 163) by Mann–Whitney U test. Overall survival was analyzed by Kaplan–Meier estimation in 297 TCGA-COAD patients with complete survival data, stratified by mean expression (high, n = 69; low, n = 228), with log-rank test and Cox regression for hazard ratios. For pathway analysis, genes were ranked per cohort by Pearson correlation with IGF2BP1 and analyzed by pre-ranked GSEA (v4.3.3, 1,000 permutations) using the MSigDB Hallmark collection (v2026.1.Hs, (24)) and curated metastasis gene sets (Rickman, Bidus, Cromer, Chandran, Jaeger; downloaded August 19, 2026). Analyses were done in R (4.5.3) and RStudio (2026.06.0+242).

**Fig. 1.**
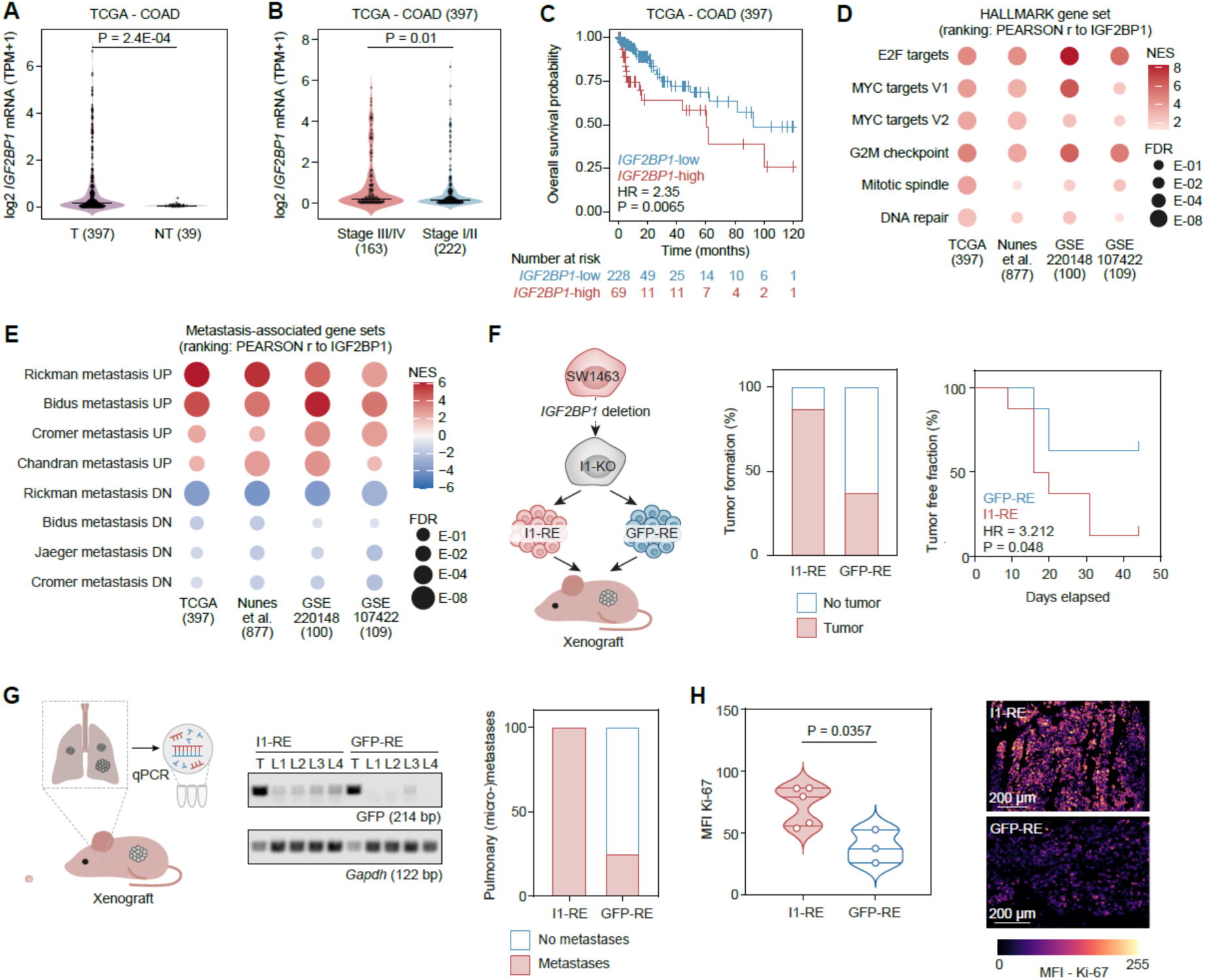
IGF2BP1 expression is associated with proliferative programs in CRC and promotes tumor formation in vivo. **(A)** IGF2BP1 mRNA expression in primary CRC (T) and non-malignant tissue (NT). **(B)** IGF2BP1 expression according to tumor stage. Data were derived from TCGA-COAD and statistical comparisons in A and B were performed using the Mann–Whitney U test. **(C)** Kaplan–Meier analysis of overall survival stratified by mean IGF2BP1 expression (log-rank test; HR from Cox regression). **(D, E)** Pre-ranked GSEA of IGF2BP1-correlated genes (Pearson correlation) across four CRC cohorts, showing selected Hallmark (D) and metastasis-associated gene sets (E). Dot color indicates normalized enrichment score (NES) and size indicates FDR. **(F)** Subcutaneous xenografts of SW1463 IGF2BP1-knockout cells re-expressing GFP-IGF2BP1 (I1-RE) or GFP (GFP-RE), injected bilaterally into four mice per group. Tumor formation frequency and tumor-free fraction over time are shown (log-rank test; HR from Cox regression). **(G)** PCR detection of GFP sequences in lung tissue, with murine Gapdh as control. T, tumor positive control; L1–L4, individual mouse lungs. Right, percentage of mice with detectable pulmonary GFP sequences. **(H)** Ki-67 fluorescence intensity in xenograft sections (unpaired two-tailed t-test) with representative images (scale bars, 200 µm). Violin plots show individual values, medians, and quartiles.

### Single-cell RNA-seq and gene set enrichment analysis

Single-cell RNA-seq data from the CRC atlas (25) were analyzed to compare IGF2BP1-positive and IGF2BP1-negative malignant cells (cluster 4). Cells with detectable IGF2BP1 expression were classified as positive and matched to IGF2BP1-negative cells according to sequencing depth, yielding 4,483 cells per group from 172 donors. Raw counts were aggregated by donor and IGF2BP1 status to generate paired pseudobulk profiles. Donors with at least ten matched cell pairs were retained for differential expression analysis (46 donors, 92 pseudobulk samples). Lowly expressed genes were filtered using the edgeR function filterByExpr. Counts were normalized using the trimmed mean of M-values (TMM) method. Differential expression was assessed using edgeR with a donor-adjusted design (∼ donor + IGF2BP1 status), quasi-likelihood dispersion estimation and robust quasi-likelihood model fitting. Hallmark gene set enrichment analysis was performed using fgsea and gene sets retrieved from MSigDB through msigdbr. Genes were ranked by the signed square root of the edgeR quasi-likelihood F-statistic, with duplicate gene symbols resolved by retaining the highest absolute ranking score. Gene sets containing 15–500 genes were tested, and Benjamini–Hochberg-adjusted P values below 0.05 were considered significant.

### Cell culture, transfection and viral transduction

Cell lines were purchased from ATCC in 2024. Cells were tested regularly for mycoplasma by PCR. SW480 (CCL-228) and HT-29 (HTB-38) cells were cultured in RPMI-1640, SW1463 (CCL-234) in DMEM/Ham’s F-12, and PANC-1 (CRL-1469) and HEK293T/17 (CRL-11268) cells in DMEM, each with 10% FBS and 1% GlutaMAX at 37 °C and 5% CO₂. IGF2BP1-knockout (I1-KO) SW480 and SW1463 cells were generated by CRISPR/Cas9 as described (16). HEK293T/R-Spondin-1 (CRL-3797) and L-Wnt-3A (CRL-2647) cells were maintained in DMEM supplemented with 10% fetal bovine serum and 1% GlutaMAX, under selection with 300 µg/ml Zeocin or 400 µg/ml G418, respectively (Thermo Fisher Scientific). For conditioned medium production, cells were incubated for 48 h in Advanced DMEM/F-12 supplemented with 1% HEPES, 1% GlutaMAX and 1% penicillin/streptomycin (Thermo Fisher Scientific).

Cells (1 × 10⁶ per 6-well) were transfected with 15 nM siRNA pools (siI1, siINCENP or non-targeting siC) using 7.5 µl Lipofectamine RNAiMAX (Thermo Fisher Scientific). For lentiviral re-expression, GFP, GFP-tagged IGF2BP1 or a KH-domain mutant of IGF2BP1 were expressed from pLVX vectors. Doxycycline-inducible expression of SBP or SBP-tagged IGF2BP1 was achieved using pLIX vectors (1 µg/ml doxycycline for 72 h). Lentiviral particles were produced in HEK293T/17 cells using the packaging plasmids psPAX2 and pMD2.G. Target cells were transduced at a multiplicity of infection (MOI) of 10. Viral titers were determined by flow cytometry (MACSQuant), and GFP-positive cells were enriched by fluorescence-activated cell sorting (FACS). BTYNB (Cayman Chemical) or DMSO was used as indicated. siRNAs, sgRNAs, plasmids, primers and antibodies are listed in Supplementary Table 1.

### Treatment, viability, proliferation and apoptosis assays

In 2D cultures, cell viability was assessed using CellTiter-Glo (Promega) after 72 h of BTYNB treatment. EC₅₀ values were determined by four-parameter nonlinear regression using GraphPad Prism (v11). For proliferation assays, 4 × 10³ cells per well were monitored for 6 days using an IncuCyte S3 live-cell imaging system (Sartorius). Cell proliferation was quantified using AI-based single-cell segmentation (IncuCyte AI Cell Health Analysis Software). Caspase-3/7 activity was measured after 72 h of BTYNB treatment using Caspase-Glo 3/7 (Promega). SW480 and SW1463 cells were treated with BTYNB (1.25–20 µM and 0.3125–5 µM, respectively) and paclitaxel (0.3125–5 nM) in 5 × 5 dose matrices for 48 h. Cell viability was assessed using CellTiter-Glo in triplicate. Drug interactions were analyzed using the Highest Single Agent (HSA) reference model implemented in SynergyFinder (https://synergyfinder.org) (26). The 2 × 2 dose region exhibiting the highest synergy score is highlighted in Fig. 6B.

For spheroid formation, 4 × 10³ cells were seeded into 96-well ultra-low attachment round-bottom plates (Corning) 24 h after transfection and centrifuged at 300 × g for 3 min. Spheroid development was monitored using a BZ-800 microscope (Keyence), Spheroid viability and caspase-3/7 activity were assessed 5 days after seeding using CellTiter-Glo and Caspase-Glo 3/7, respectively (Promega).

### Patient-derived organoid culture and drug treatment

CRC organoids (CRC2, CRC7, CRC50, CRC51, CRC52, CRC260) were obtained from ATCC (catalog numbers in Supplementary Table 2), with ethical approval and consent provided by the supplier.

Organoids were embedded in 50–100 µl Matrigel (Corning) domes and maintained with weekly passaging in complete organoid medium consisting of Advanced DMEM/F-12 supplemented with 10% R-Spondin-1- and 10% Wnt-3A-conditioned medium, 1% HEPES, 1% GlutaMAX, 1% penicillin/streptomycin, 1× B27, 1× N-2, 10 mM nicotinamide, 1.25 mM N-acetylcysteine, 10 µM forskolin, 10 ng/ml FGF-10, 10 ng/ml Noggin, 5 ng/ml EGF, 5 µM Y-27632, 500 nM SB431542, 200 nM hydrocortisone, 10 nM gastrin, 10 nM prostaglandin E2 and 0.2% Primocin. Drug assays were performed between passages 5 and 10 in reduced organoid medium, prepared by omitting forskolin, FGF-10, EGF, hydrocortisone, gastrin and prostaglandin E2 from the complete formulation. For these assays, organoids were harvested in ice-cold organoid medium, centrifuged at 300 × g for 5 min at 4°C and seeded in 5 µl Matrigel domes in 96-well plates. After 48 h, organoids were treated with the indicated compounds and concentrations. Organoid growth was monitored for 72 h using an IncuCyte S3 system (Sartorius) with the organoid analysis module. Growth was quantified as segmented organoid area normalized to the area at treatment initiation. Experiments were performed in biological triplicates. Organoid viability was assessed using CellTiter-Glo (Promega), and EC₅₀ values were determined by four-parameter nonlinear regression. For combination treatments, organoids were exposed to the indicated concentrations of BTYNB and paclitaxel, either individually or in combination, for 72 h. Drug interactions were assessed using ordinary two-way ANOVA with BTYNB and paclitaxel as independent factors. The BTYNB × paclitaxel interaction term was used to evaluate deviations from additivity, and Šídák’s multiple comparisons test was applied for selected pairwise comparisons. To quantify the magnitude and direction of the interaction, an inverted interaction contrast was calculated as Δ = vBTYNB + vPaclitaxel − vCombination − vDMSO, where v denotes DMSO-normalized fractional viability. Positive values indicate a greater reduction in viability than expected from additive single-agent effects. Drug interactions were additionally evaluated using the Highest Single Agent (HSA) and Bliss independence models. For each independent experiment, synergy scores were calculated as HSA = min(vBTYNB, vPaclitaxel) − vCombination and Bliss = (vBTYNB × vPaclitaxel) − vCombination. Positive scores indicate synergy relative to the respective reference model. HSA and Bliss scores from three independent experiments per organoid line were compared against zero using two-tailed one-sample t-tests, with Holm correction for multiple testing across the six organoid lines separately for each model.

For morphological analysis, BTYNB-treated organoids were fixed in 4% paraformaldehyde for 30 min at room temperature, stained with Hoechst and TRITC-conjugated phalloidin, and imaged using a Leica Viventis SCAPE light-sheet microscope with standardized acquisition settings.

### Xenograft experiments

Female six-week-old, athymic nude mice (FOXN1^nu/nu^; Charles River) were used in accordance with the guidelines of Martin Luther University Halle-Wittenberg and approved by the state administration office of Saxony-Anhalt (42502-2-1625 MLU). SW1463 I1-KO cells re-expressing GFP-IGF2BP1 (I1-RE) or GFP (GFP-RE) were injected bilaterally into the flanks of four animals per group (3 × 10⁶ cells per flank in 50% v/v Matrigel; n = 8 tumors per group). Tumors were measured by caliper every 3–4 days for 45 days, and none exceeded 1.5 cm in diameter. Tumors and lungs were harvested at the end.

Fresh-frozen tumor sections were stained for Ki-67 with the Opal 7 system (Akoya Biosciences), imaged on a Leica TCS SP8X confocal microscope, and the mean fluorescence intensity was quantified in QuPath (v0.7.0). Lungs were homogenized in TRIzol to extract total RNA; cDNA was synthesized with M-MLV reverse transcriptase (Promega) and random hexamers. PCR (30 cycles) with primers against GFP (214 bp) and murine Gapdh (122 bp) was resolved on ethidium-bromide-stained agarose gels to detect pulmonary tumor cells.

### Western blotting, immunofluorescence and cell cycle analysis

Western blotting was performed as described (17), with vinculin as loading control. For immunofluorescence, cells on coverslips were fixed in ice-cold methanol (10 min, −20 °C), blocked in 1% BSA/PBS, incubated with anti-α-tubulin (1 h) and a fluorescent secondary antibody (1 h), counterstained with DAPI (1:50,000) and mounted. Images were acquired on a Leica TCS SP8X confocal microscope, and at least 50 mitotic figures per condition were scored for spindle abnormalities.

For cell cycle analysis, cells were harvested 72 h after transfection or 48 h after BTYNB treatment and fixed in 70% ethanol (−20 °C, overnight). DNA was stained with propidium iodide (1:1000; Miltenyi Biotec; 30 min, 37 °C) in PBS with RNase A (2 µg/ml; Sigma-Aldrich). DNA content was measured on a MACSQuant Analyzer and analyzed in FlowJo (v10.8.1) using the Watson (pragmatic) fit to determine G1, S, G2M and >4n fractions.

### RNA immunoprecipitation and mRNA decay analysis

RNA immunoprecipitation (RIP) and RT-qPCR were performed as previously described (15). RIP was conducted in SW480 IGF2BP1-KO cells re-expressing GFP-IGF2BP1 (GFP-I1-RE) or GFP (GFP-RE). Where indicated, GFP-I1-RE cells were treated with 5 µM BTYNB or DMSO for 24 h. RNA was isolated using TRIzol, and RIP enrichment was quantified by RT-qPCR using H2AC6 and H3C15 for normalization.

For mRNA decay analysis, SW480 cells were pretreated with 5 µM BTYNB or DMSO for 24 h, followed by transcriptional inhibition with 5 µM actinomycin D (Sigma-Aldrich). RNA was collected at the indicated time points, and transcript abundance was determined by RT-qPCR. Relative expression was calculated using the ΔΔCt method with RPLP0, EEF2 and GAPDH as reference genes. Decay curves were normalized to the respective time point 0, log₂-transformed, and fitted by linear regression. Transcript half-lives (t½) were calculated from the fitted slopes. Statistical differences between DMSO- and BTYNB-treated decay curves were assessed using the modified chi-square method for comparing arbitrary curves described by Hristova and Wimley (27).

### RNA sequencing and differential expression analysis

Total RNA was polyA-enriched and sequenced by Novogene (München) on an Illumina NovaSeq platform (three biological replicates per condition): SW480 siI1 vs. siC, HT-29 SBP-I1 vs. SBP, and SW480 BTYNB (5 µM, 72 h) vs. DMSO. Reads were trimmed with Cutadapt 4.4, aligned to GRCh38 and counted with featureCounts 2.0.6. Differential expression was determined with edgeR 4.10.1 (TMM normalization, (28)); genes with FDR < 0.05 were considered significant. For GSEA, genes were ranked by log₂ fold change and analyzed in GSEA 4.3.3 with the Hallmark and GO-BP collections.

Genes downregulated in both the BTYNB and siI1 datasets (FDR < 0.05; n = 115) were analyzed for over-representation of GO-BP terms with clusterProfiler (29) and org.Hs.eg.db (all genes tested in both contrasts; Benjamini–Hochberg-adjusted P < 0.05). Redundant terms were reduced with the simplify function (cutoff 0.6), and the ten most significant terms are shown in Fig. 4A. For the recurrence analysis (Fig. 4B), simplified terms were filtered by keywords for mitosis, chromosome segregation and cell cycle (seven terms), and genes were ranked by the number of these terms in which they are annotated. The ten top genes are shown, with the four CPC genes always included.

### eCLIP data and multi-model IGF2BP1 depletion analysis

Public IGF2BP1 eCLIP peak files from H9 embryonic stem cells (GSM2071742, GSM2071745; GSE78507/GSE78509) (30) and ENCODE HepG2 (ENCFF486BXN, ENCFF976DBP) and K562 cells (ENCFF435MEM, ENCFF701YCW) (31) were integrated in R. Published peak calls were used without an additional signal or P-value cutoff. BED starts were converted to 1-based coordinates, and hg19 peaks were lifted to hg38 with the UCSC chain file (rtracklayer; one-to-one mappings only). Replicate peaks were merged within each cell type, and consensus regions supported by at least two of three cell types were defined with ChIPpeakAnno (findOverlapsOfPeaks, maxgap = 20, connectedPeaks = “min”). Peaks were assigned to one transcript per gene (longest CDS, TxDb.Hsapiens.UCSC.hg38.knownGene) and classified as 5ʹUTR, CDS, 3ʹUTR or intron by their midpoint. The CLIP viewer plots strand-oriented peak midpoints on this transcript model for user-defined gene lists, with exon-only, high-confidence or whole-gene display modes. Lollipop height is the mean log2 enrichment, dot size the number of supporting cell types, and color the mean P across all six experiments (non-detection = 0), a descriptive confidence score rather than a combined P value. H9 scores were not aggregated because their scale differs from ENCODE.

The DEG viewer summarizes gene-level log₂ fold changes and P values from 15 siRNA-mediated IGF2BP1-depletion datasets (8305C, C643, Hth74, MB-1, A-549, BE2C, KELLY, CFPAC, PANC-1, PANC0327 (GSE319148), ES-2, HepG2, HUH-7, MV-3, SW480_CRC). For each gene, the dominant direction, the number of agreeing models and the mean log₂ fold change of those models were calculated. P values from agreeing models were combined by Fisher’s method (log scale; Benjamini–Hochberg-adjusted), and directional agreement was assessed by a one-sided binomial test. Codirectional summaries describe the recurrence of expression changes across models. Bubbles show the codirectional mean log₂ fold change, with size indicating agreement (14–15, 12–13, 10–11 or 8–9 of 15 models) and color −log₁₀ of the adjusted combined P value, capped at the 98th percentile. eCLIP binding information was integrated for annotation without restricting the analysis to genes with detectable IGF2BP1 binding.

### Label-free quantification mass spectrometry

PANC-1 cells were transfected with siC or si_I1 (n = 3 per condition) and harvested after 72 h. Proteins were digested by a modified FASP protocol (32), and peptides were analyzed with an Ultimate 3000 RSLC nano-HPLC system coupled to a timsTOF Ultra-2 mass spectrometer (Bruker Daltonik) in DIA-PASEF mode (*m/z* 100–1700; 0.64–1.45 V·s/cm²; 24 windows *m/z* 400–1000; collision energies 20–59 eV). LC separation was performed using a 60-min gradient (3–30% acetonitrile, 0.1% formic acid, 300 nL/min) on a self-packed C18 column (75 µm × 250 mm; ReproSil-Pur 1.9 µm). Data were searched with directDIA in Spectronaut (v19; Biognosys) and Proteoscape against the UniProt human proteome database (February 2025; 20,427 entries) with default thresholds (precursor and protein q-values ≤ 0.01; at least two peptides per protein; up to two missed cleavages). Methionine oxidation and N-terminal acetylation were set as variable and cysteine carbamidomethylation as fixed modifications. Only proteotypic peptides were quantified, and only proteins quantified in at least two of three replicates per condition were considered. Differential abundance was assessed by permutation-based FDR.

### Statistical analysis

Unless stated otherwise, experiments were performed in at least three independent biological replicates, and data are shown as mean ± SD. Statistical analyses were performed using GraphPad Prism or R, as appropriate. Statistical tests, sample sizes, and multiple testing corrections are specified in the respective Methods sections and figure legends. Unless otherwise indicated, P < 0.05 was considered statistically significant.

## Results

### IGF2BP1 expression is associated with proliferative programs in CRC and promotes tumor formation *in vivo*

We first investigated the clinical relevance of IGF2BP1 in CRC using the TCGA-COAD cohort. IGF2BP1 mRNA expression was significantly elevated in primary tumors compared with non-malignant tissue (Fig. 1A). Moreover, expression increased with tumor stage (Fig. 1B), and patients with above-mean IGF2BP1 expression showed significantly reduced overall survival (Fig. 1C). Stratification by consensus molecular subtype (CMS) further revealed elevated IGF2BP1 expression in CMS2 and CMS4 tumors (Supplementary Fig. S1A), which are characterized by distinct proliferative and mesenchymal transcriptional programs, respectively (7).

Given the established role of IGF2BP1 in sustaining oncogenic proliferation in neuroblastoma, ovarian cancer, and other malignancies, we next investigated whether similar transcriptional programs accompany IGF2BP1 expression in CRC. Pre-ranked gene set enrichment analysis (GSEA) of IGF2BP1-correlated genes across four independent CRC cohorts (TCGA-COAD, GSE220148, GSE107422, and E-MTAB-12862) consistently revealed positive enrichment of proliferation-associated Hallmark gene sets, including E2F and MYC targets, G2/M checkpoint, mitotic spindle, and DNA repair (Fig. 1D). These findings linked IGF2BP1 expression to cell cycle progression and mitotic regulation, consistent with the proliferative characteristics of CMS2 tumors. In addition, metastasis-associated gene sets showed consistent enrichment patterns across the four cohorts, with positive enrichment of metastasis-upregulated signatures and negative enrichment of metastasis-downregulated signatures (Fig. 1E), compatible with the invasive and mesenchymal features characteristic of CMS4.

To determine whether IGF2BP1 functionally contributes to tumor formation, we employed SW1463 IGF2BP1-knockout cells re-expressing GFP-IGF2BP1 (I1-RE) or GFP alone (GFP-RE), with knock-out and re-expression confirmed by Western blotting (Supplementary Fig. S1B). Following subcutaneous injection into immunodeficient mice, I1-RE cells formed palpable tumors more frequently and earlier than GFP-RE cells (Fig. 1F). Interestingly, pulmonary GFP sequences were detected in all I1-RE mice but only one of four GFP-RE mice (Fig. 1G), consistent with enhanced pulmonary dissemination and with IGF2BP1-associated metastatic transcriptional programs in CRC and neuroblastoma (33, 34). Importantly, Ki-67 staining revealed significantly increased proliferative activity in I1-RE tumors (Fig. 1H and Supplementary Fig. S1C, D), providing functional support for the association between IGF2BP1 and proliferative transcriptional programs observed in CRC patients.

### IGF2BP1 promotes proliferation and survival of CRC cells

To assess the association between IGF2BP1 expression and proliferative programs in CRC cells, we analyzed a single-cell CRC atlas ((GSE285117) (25)). We compared gene expression between sequencing-depth-matched IGF2BP1-positive and -negative cancer cells and performed GSEA. We found strong enrichment of E2F and MYC targets, G2/M checkpoint, and mitotic spindle signatures in IGF2BP1-positive cells, whereas metabolic and inflammatory response programs were predominantly depleted (Fig. 2A, B).

**Fig. 2.**
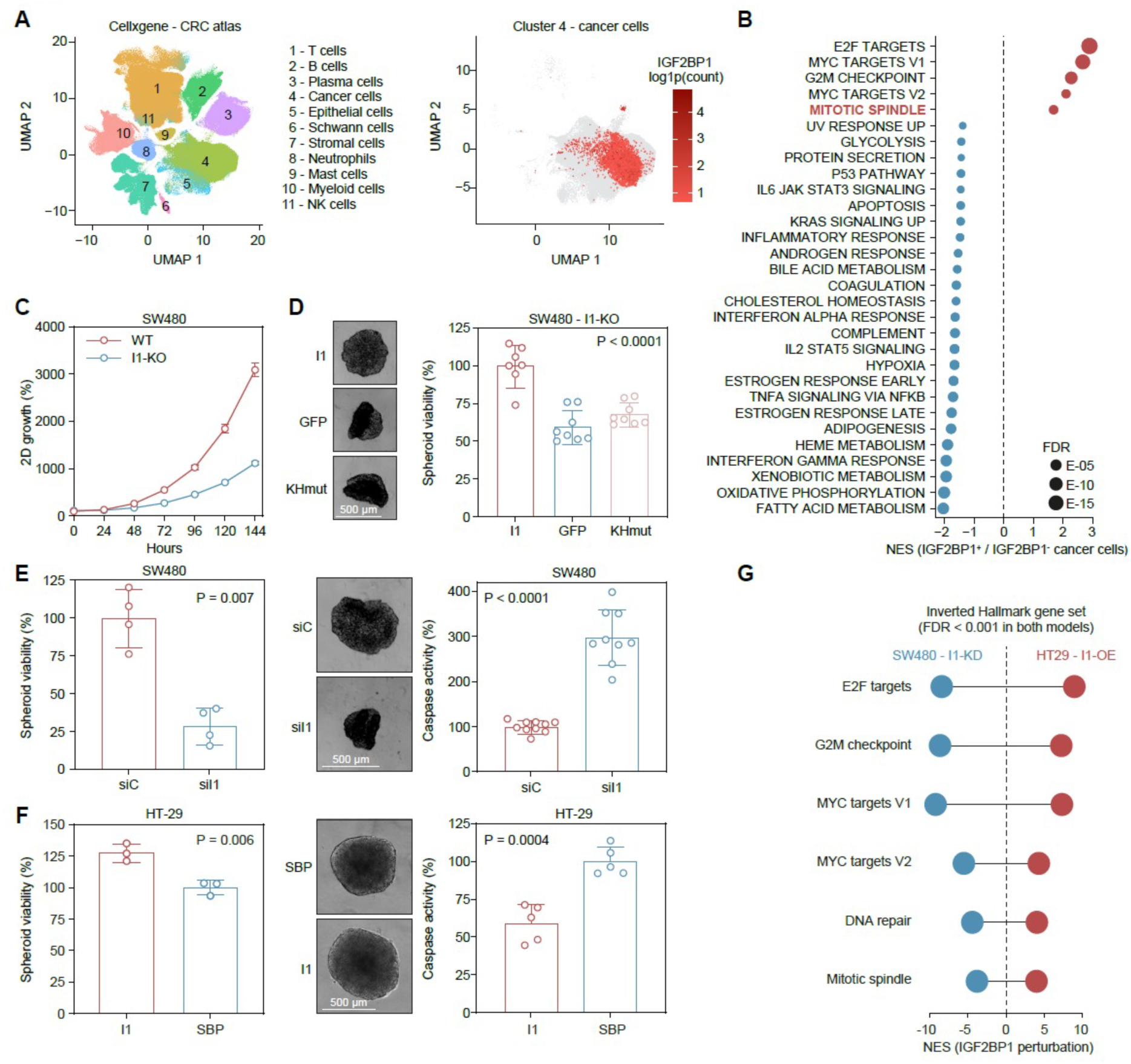
IGF2BP1 promotes proliferation and survival of CRC cells. **(A)** UMAP of the CRC single-cell atlas (GSE285117) showing annotated cell populations and IGF2BP1 expression in malignant cells (cluster 4). **(B)** Pre-ranked Hallmark GSEA comparing sequencing-depth-matched IGF2BP1-positive and - negative cancer cells (n = 4,483 cells per group). Dot position indicates normalized enrichment score (NES) and size indicates false discovery rate (FDR). **(C)** Live-cell imaging of SW480 wild-type (WT) and IGF2BP1-knockout (I1-KO) cells over 144 h. **(D)** Spheroid viability of SW480 I1-KO cells re-expressing GFP-IGF2BP1 (I1), GFP, or an IGF2BP1 KH1-4-domain mutant (KHmut). **(E, F)** Spheroid viability and caspase activity following IGF2BP1 siRNA depletion in SW480 cells (E) or doxycycline-induced SBP-IGF2BP1 expression in HT-29 cells (F), with respective controls. Viability and caspase activity were assessed using CellTiter-Glo and Caspase-Glo 3/7, respectively, five days after seeding. **(G)** Hallmark GSEA upon IGF2BP1 knockdown in SW480 or overexpression in HT-29 cells, showing reciprocally regulated gene sets (FDR < 0.001 in both models). Data in C–F are mean ± SD; individual points represent independent experiments. Statistical significance was assessed by one-way ANOVA with Tukey’s multiple comparisons test (D) or unpaired two-tailed t-test (E, F). Representative spheroid images are shown in D–F (scale bars, 500 µm).

To investigate the functional contribution of IGF2BP1, we employed genetic loss-, rescue-, and gain-of-function approaches (Supplementary Fig. S2A). IGF2BP1 knockout markedly impaired proliferation of SW480 cells (Fig. 2C). Re-expression of GFP-IGF2BP1 restored viability in 3D spheroids, whereas GFP alone or an RNA-binding-deficient KH-domain mutant failed to rescue this phenotype (Fig. 2D, Supplementary Fig. S2B). Similarly, transient IGF2BP1 depletion reduced spheroid viability and increased caspase activity (Fig. 2E, Supplementary Fig. S2B). Conversely, inducible IGF2BP1 expression in HT-29 cells, which express little endogenous IGF2BP1, enhanced spheroid viability and suppressed caspase activity (Fig. 2F). Together, these findings demonstrate that IGF2BP1 promotes CRC cell growth and survival in an RNA-binding-dependent manner.

To investigate the transcriptional changes associated with these proliferative programs, we performed RNA-sequencing following IGF2BP1 knockdown in SW480 cells and inducible overexpression in HT-29 cells. GSEA revealed reciprocal regulation of E2F and MYC targets, G2/M checkpoint, DNA repair, and mitotic spindle signatures, which were negatively enriched upon IGF2BP1 depletion and positively enriched following its overexpression (Fig. 2G). While the enrichment of E2F and MYC target signatures is consistent with established IGF2BP1-dependent regulation of E2F1 and MYC/MYCN (13, 15, 16) the pronounced changes in mitotic programs suggested additional regulatory mechanisms underlying IGF2BP1-driven proliferation.

### BTYNB impairs CRC cell viability, induces apoptosis and causes mitotic defects

To test the consequences of pharmacological IGF2BP1 inhibition, we treated SW480 cells with BTYNB, a small-molecule inhibitor that interferes with IGF2BP1 binding to target mRNAs. BTYNB reduced cell viability in a dose-dependent manner (EC₅₀ = 2.97 ± 0.41 µM). At higher cell densities, 5 µM BTYNB reduced viability by approximately 50% and increased caspase activity (Fig. 3A). Interestingly, treated cells became enlarged and flattened, with pronounced alterations in nuclear morphology (Fig. 3B), suggesting defects in cell division beyond the established regulation of E2F/MYC-associated proliferative programs.

**Fig. 3.**
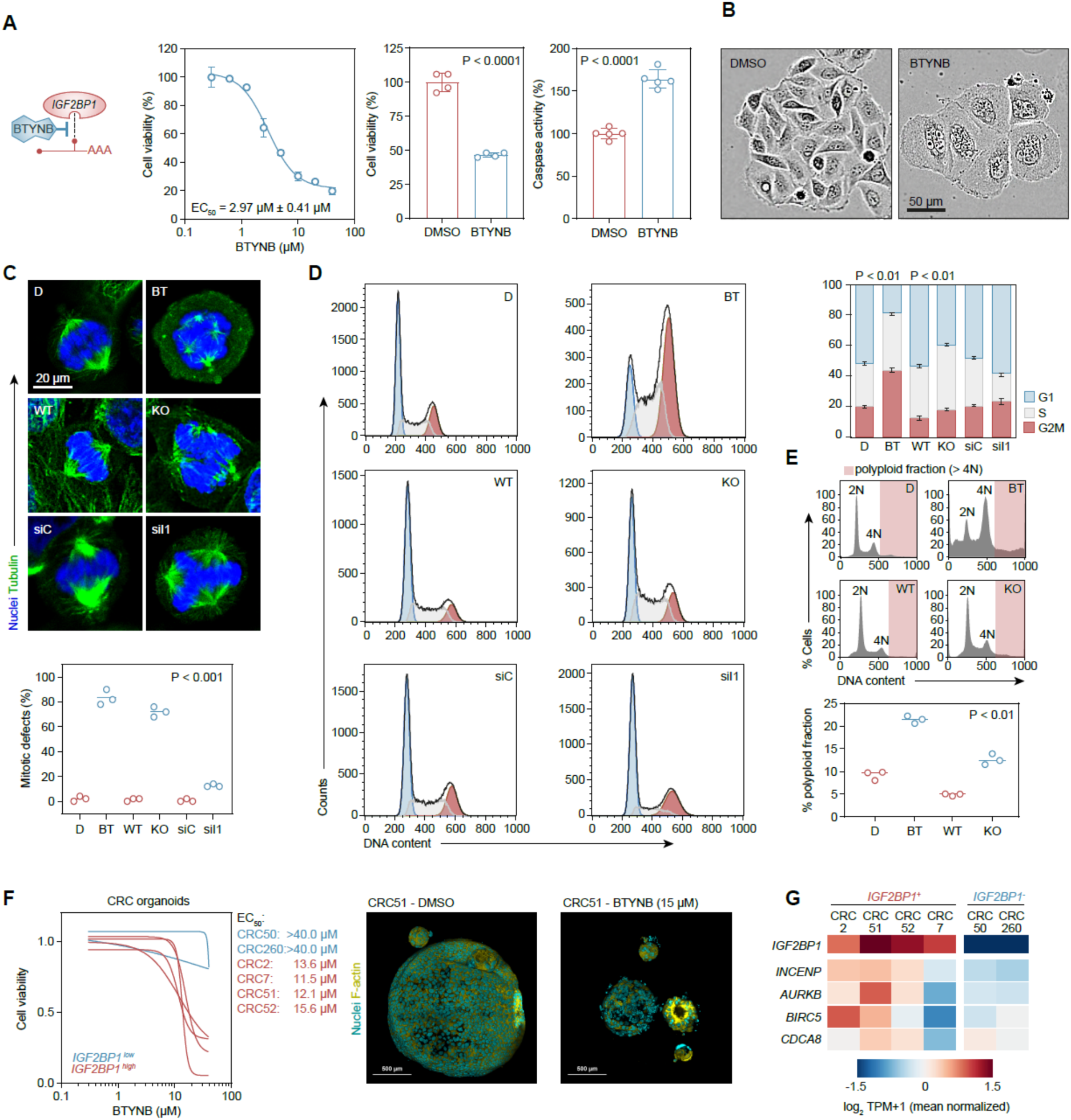
BTYNB impairs CRC cell viability, induces mitotic defects and preferentially affects IGF2BP1-expressing organoids. **(A)** Schematic of BTYNB-mediated inhibition of IGF2BP1–RNA interactions, dose-dependent SW480 cell viability, and viability and caspase activity following treatment with 5 µM BTYNB or DMSO for 72 h. **(B)** Representative brightfield images of treated SW480 cells (scale bar, 50 µm). **(C)** Representative α-tubulin (green) and nuclear (blue) staining of mitotic SW480 cells following BTYNB treatment, IGF2BP1 knockout (KO), or siRNA depletion (siI1), with respective controls. Quantification shows mitotic spindle defects (≥50 mitotic figures per condition; scale bar, 20 µm). **(D)** Cell cycle analysis by PI flow cytometry using the Watson pragmatic model in FlowJo, showing fitted G1, S, and G2/M distributions and their quantification. P values indicate G2/M comparisons (DMSO vs. BTYNB; WT vs. KO). **(E)** Representative PI fluorescence histograms with gating and quantification of cells with >4N DNA content (shaded area). **(F)** BTYNB dose–response curves and EC₅₀ values in six patient-derived CRC organoid lines (72 h), with representative images of CRC51 organoids (nuclei, blue; F-actin, yellow; scale bars, 500 µm). EC₅₀ values for CRC50 and CRC260 exceeded 40 µM, the highest tested concentration. **(G)** Mean-normalized log₂ (TPM + 1) expression of IGF2BP1 and chromosomal passenger complex (CPC) components in CRC organoids. Data in A and C–E are mean ± SD or individual values; each point represents an independent experiment. P values were determined by unpaired two-tailed t-tests for pairwise comparisons.

To investigate this further, we examined spindle organization by α-tubulin and DNA staining. BTYNB markedly increased the proportion of mitotic cells exhibiting spindle defects (83.3% vs. 2% in DMSO). Similar abnormalities were observed in IGF2BP1-knockout cells (72% vs. 1.3% in WT) and, to a lesser extent, following transient IGF2BP1 depletion (12.7% vs. 0.7% in siC; Fig. 3C). DNA-content analysis further revealed a pronounced increase in the G2/M fraction following BTYNB treatment (43% vs. 19% in DMSO), accompanied by an increased proportion of cells with DNA content exceeding 4N (21% vs. 9%; Fig. 3D, E). IGF2BP1-knockout cells showed a similar, although less pronounced, increase in G2/M and >4N populations (12% vs. 5% for >4N; Fig. 3D, E). In contrast, transient IGF2BP1 depletion predominantly increased the G1 fraction and reduced S-phase representation, with a modest, non-significant increase in G2/M (Fig. 3D). This is consistent with the previously described role of IGF2BP1 in E2F-dependent G1/S progression (15). Together, these findings indicate that IGF2BP1 contributes to mitotic integrity in CRC cells in addition to its established role in G1/S progression.

We next examined whether BTYNB sensitivity extends to patient-derived CRC organoids. Four organoid lines with higher IGF2BP1 expression (CRC2, CRC7, CRC51 and CRC52) showed pronounced dose-dependent reductions in viability (EC₅₀ = 11.5–15.6 µM). BTYNB treatment also disrupted organoid architecture, as illustrated by CRC51 (Fig. 3F). In contrast, CRC50 and CRC260, which expressed substantially less IGF2BP1 (Fig. 3G), retained approximately 80% viability even at the highest tested concentration (40 µM), precluding reliable EC₅₀ determination. Thus, BTYNB sensitivity was substantially associated with higher IGF2BP1 expression across the patient-derived CRC organoid models examined.

Given the pronounced mitotic defects following IGF2BP1 perturbation, we next focused on the chromosomal passenger complex (CPC), a key regulator of chromosome segregation and cytokinesis. Expression profiling of patient-derived CRC organoids revealed variable expression of CPC components, with INCENP showing the strongest association with IGF2BP1 (Fig. 3G). We therefore investigated whether IGF2BP1 contributes to CPC regulation.

### IGF2BP1 inhibition and depletion converge on the chromosomal passenger complex

To identify molecular mechanisms underlying the mitotic defects observed upon IGF2BP1 perturbation, we compared transcriptomic changes following BTYNB treatment and siRNA-mediated IGF2BP1 depletion in SW480 cells. Despite the markedly different severity of the mitotic phenotypes observed with these two approaches (Fig. 3C–E), we identified 115 genes downregulated under both conditions (FDR < 0.05). These genes were enriched for biological processes related to mitotic cell cycle progression, chromosome segregation, cytokinesis, DNA replication, and repair (Fig. 4A). To prioritize candidate effectors, we ranked these genes by their representation across seven mitosis-associated Gene Ontology biological process (GO-BP) terms. Remarkably, all four components of the chromosomal passenger complex (CPC)—INCENP, AURKB, CDCA8, and BIRC5—ranked among the most frequently represented genes, each occurring in five of the seven terms (Fig. 4B). The CPC coordinates chromosome segregation and cytokinesis through its dynamic localization during mitosis, as illustrated schematically in Fig. 4B. The coordinated representation of all four components therefore prompted us to investigate their regulation by IGF2BP1.

**Fig. 4.**
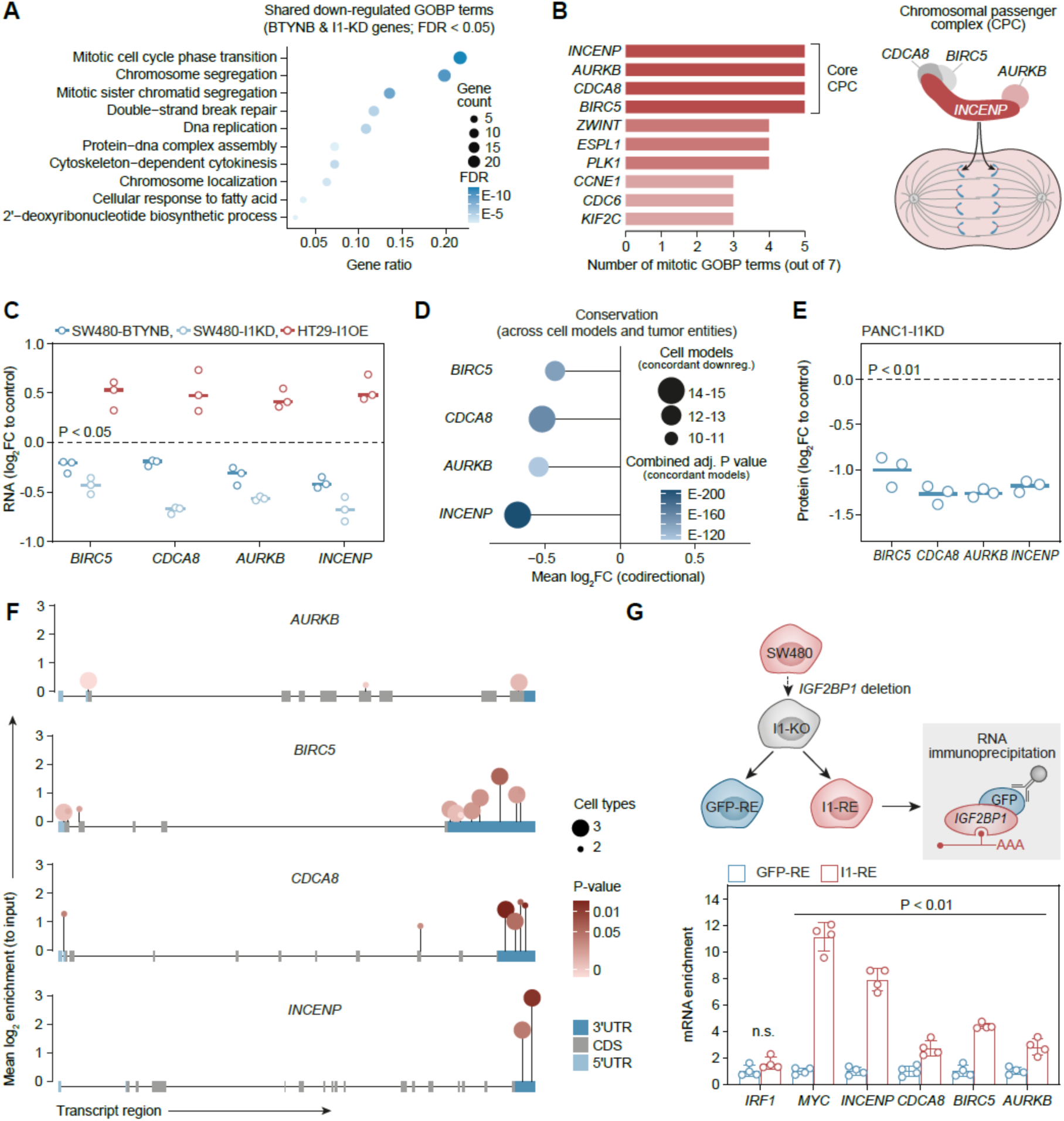
IGF2BP1 regulates chromosomal passenger complex expression and associates with CPC-encoding mRNAs. **(A)** GO biological process (GO-BP) enrichment of 115 genes downregulated following both BTYNB treatment and IGF2BP1 knockdown in SW480 cells (FDR < 0.05). Dot size indicates gene count and color indicates FDR. **(B)** Ten genes most frequently represented among seven mitotic GO-BP terms (left) and schematic of the chromosomal passenger complex (CPC) during mitosis (right). **(C)** CPC mRNA expression changes following BTYNB treatment (5 µM, 72 h) or IGF2BP1 knockdown in SW480 cells and IGF2BP1 overexpression in HT-29 cells (n = 3 biological replicates; P < 0.05). **(D)** Codirectional mean log₂ fold changes following IGF2BP1 depletion across cancer cell models and tumor entities. Dot size indicates the number of models with concordant downregulation and color indicates the adjusted combined P value calculated from these models. **(E)** CPC protein abundance changes measured by label-free mass spectrometry in PANC-1 cells following IGF2BP1 knockdown (72 h; n = 3; P < 0.01). **(F)** IGF2BP1 eCLIP binding profiles across CPC transcripts. Lollipops indicate mean log₂ enrichment over input, with dot size representing the number of supporting cell types (two or three) and color indicating P values. Transcript regions are annotated as 5ʹUTR, CDS, and 3ʹUTR. **(G)** GFP-RNA immunoprecipitation in SW480 IGF2BP1-knockout cells re-expressing GFP-IGF2BP1 (I1-RE) or GFP (GFP-RE). mRNA enrichment is shown relative to GFP-RE; MYC and IRF1 serve as positive and negative controls, respectively. Data in C, E, and G show individual biological replicates and means (± SD in G). P values were < 0.01 for all comparisons in G except IRF1 (n.s.).

Consistent with this hypothesis, all four CPC transcripts were downregulated following BTYNB treatment or IGF2BP1 knockdown in SW480 cells and upregulated upon IGF2BP1 overexpression in HT-29 cells (Fig. 4C). To assess whether this regulation extends beyond CRC, we integrated RNA-seq data from 15 cancer cell models representing a variety of tumor entities following IGF2BP1 depletion. INCENP and CDCA8 were downregulated in all 15 models, whereas AURKB and BIRC5 decreased in 13 of 15, indicating a broadly conserved response across tumor entities (Fig. 4D). In agreement with these transcriptional changes, label-free quantitative mass spectrometry in the independent PDAC-derived PANC-1 model revealed an approximately twofold reduction in all four CPC proteins following IGF2BP1 knockdown (Fig. 4E). Together, these findings identify CPC expression as a recurrent downstream response to IGF2BP1 perturbation at both the mRNA and protein levels.

We next investigated whether IGF2BP1 directly associates with CPC-encoding transcripts. Analysis of IGF2BP1 eCLIP datasets from H9 embryonic stem cells, HepG2, and K562 cells identified binding sites in all four CPC mRNAs (Fig. 4F). Prominent binding was observed within the 3ʹUTRs of INCENP, CDCA8, and BIRC5, whereas AURKB showed weaker binding outside the 3ʹUTR. To validate these interactions in CRC cells, we performed RNA immunoprecipitation using SW480 IGF2BP1-knockout cells re-expressing GFP-IGF2BP1 (I1-RE) or GFP alone (GFP-RE). GFP-IGF2BP1 significantly enriched all four CPC transcripts, with INCENP showing the strongest enrichment, approaching that of the established IGF2BP1 target MYC. In contrast, the negative control IRF1 showed no significant enrichment (Fig. 4G). Together, these findings identify all four CPC transcripts as IGF2BP1-associated RNAs in CRC cells, prompting us to investigate whether pharmacological inhibition disrupts these interactions.

### BTYNB disrupts IGF2BP1–CPC mRNA interactions and destabilizes INCENP, whose depletion recapitulates mitotic defects

To test the impact of BTYNB on IGF2BP1–CPC mRNA interactions, we performed RNA immunoprecipitation in SW480 GFP-I1-RE cells following BTYNB treatment (Supplementary Fig. S3A). BTYNB markedly reduced IGF2BP1 association with INCENP, CDCA8, and BIRC5 mRNAs, as well as the established target MYC, whereas AURKB binding remained largely unaffected (Fig. 5A). Since IGF2BP1 is known to promote target mRNA stability, we next examined the consequences of BTYNB treatment for mRNA decay. Following 24 h of BTYNB pretreatment, actinomycin D chase experiments revealed significantly different decay curves for both MYC and INCENP, with accelerated transcript decay under BTYNB treatment. The estimated mRNA half-lives decreased from 0.55 to 0.33 h for MYC and from 8.48 to 5.51 h for INCENP (Fig. 5B). These findings indicate that BTYNB disrupts IGF2BP1 association with selected CPC transcripts and compromises INCENP mRNA stability.

**Fig. 5.**
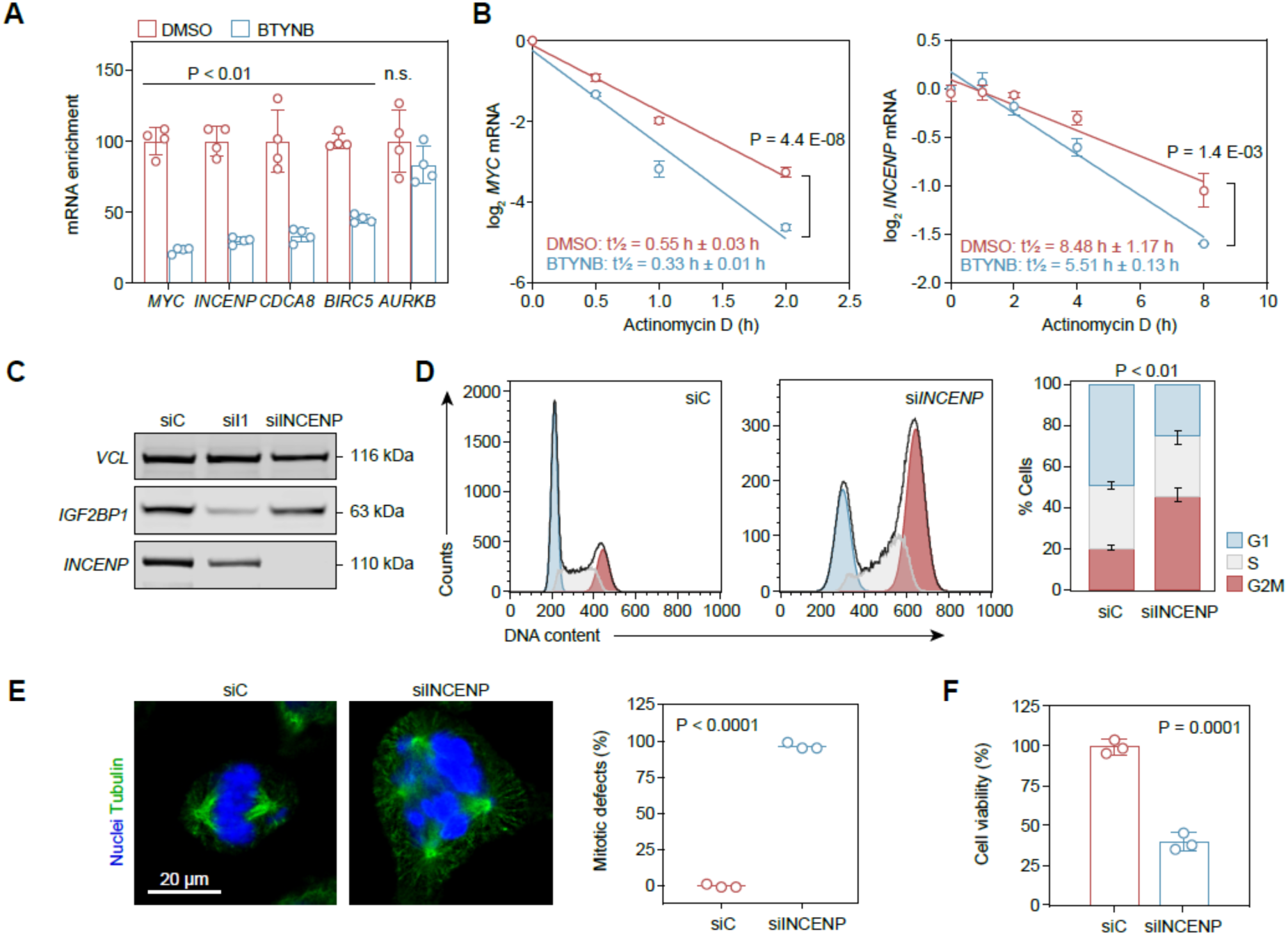
BTYNB disrupts IGF2BP1–CPC mRNA interactions and destabilizes INCENP, whose depletion recapitulates mitotic defects. **(A)** RNA immunoprecipitation using GFP antibodies in SW480 IGF2BP1-knockout cells re-expressing GFP-IGF2BP1 (GFP-I1-RE), treated with BTYNB (5 µM) or DMSO for 24 h. Relative enrichment of the indicated transcripts is shown as a percentage of DMSO controls (100%). MYC serves as a positive control. **(B)** Actinomycin D chase analysis of MYC (left) and INCENP (right) mRNA decay in SW480 wild-type cells following 24 h pretreatment with BTYNB (5 µM) or DMSO. Log₂-transformed mRNA levels normalized to time zero are shown with fitted decay curves and estimated half-lives (t½). Differences between decay curves were assessed using the modified chi-square method described by Hristova and Wimley (27). **(C)** Western blot analysis of IGF2BP1 and INCENP in SW480 cells transfected with control siRNA (siC), IGF2BP1-targeting siRNA (siI1), or INCENP-targeting siRNA (siINCENP). Vinculin (VCL) serves as a loading control. **(D)** Cell cycle analysis of SW480 cells following siC or siINCENP transfection using propidium iodide staining and flow cytometry. Representative fitted cell cycle distributions and quantification of G1, S, and G2/M fractions are shown. **(E)** Representative immunofluorescence images of α-tubulin (green) and nuclei (blue) in mitotic SW480 cells following siC or siINCENP transfection (left), and quantification of mitotic cells exhibiting spindle defects (right). Scale bar, 20 µm. **(F)** Cell viability of SW480 cells following siC or siINCENP transfection. Data represent independent experiments (n = 3–4) and are presented as mean ± SD with individual values where indicated. Except for the curve comparisons in (B), statistical significance was assessed using unpaired two-tailed Student’s t-tests. P values are indicated; n.s., not significant.

Given its central role as the CPC scaffold and its prominent association with IGF2BP1, we next investigated the functional consequences of INCENP depletion. Western blotting confirmed efficient INCENP knockdown and revealed reduced INCENP protein abundance following IGF2BP1 depletion, whereas INCENP knockdown did not affect IGF2BP1 expression (Fig. 5C). Strikingly, INCENP depletion caused pronounced G2/M accumulation (Fig. 5D) and spindle abnormalities in nearly all mitotic cells (Fig. 5E), closely resembling the defects observed following BTYNB treatment and IGF2BP1 knockout. These mitotic abnormalities were accompanied by a marked reduction in cell viability (Fig. 5F). Together, these findings identify INCENP as a functionally relevant downstream component of the IGF2BP1–CPC regulatory axis, linking impaired CPC expression to the mitotic defects observed upon IGF2BP1 perturbation.

### BTYNB synergizes with paclitaxel in CRC cells and IGF2BP1-positive organoids

Since IGF2BP1 inhibition disrupts CPC expression and causes mitotic defects, we reasoned that it might sensitize CRC cells to the microtubule-targeting drug paclitaxel (Fig. 6A). To test this, we performed 5 × 5 dose-matrix experiments in SW480 and SW1463 cells and evaluated drug interactions using the Highest Single Agent (HSA) model. Both cell lines showed concentration-dependent regions of synergy, with mean HSA δ-scores of 6.10 and 11.88, respectively. The strongest synergistic responses occurred at 2.5–5 µM BTYNB combined with 1.25–2.5 nM paclitaxel in SW480 cells and at 0.625–1.25 µM BTYNB combined with 0.625–1.25 nM paclitaxel in SW1463 cells, reaching maximum δ-scores of 22.22 and 41.09, respectively (Fig. 6B).

**Fig. 6.**
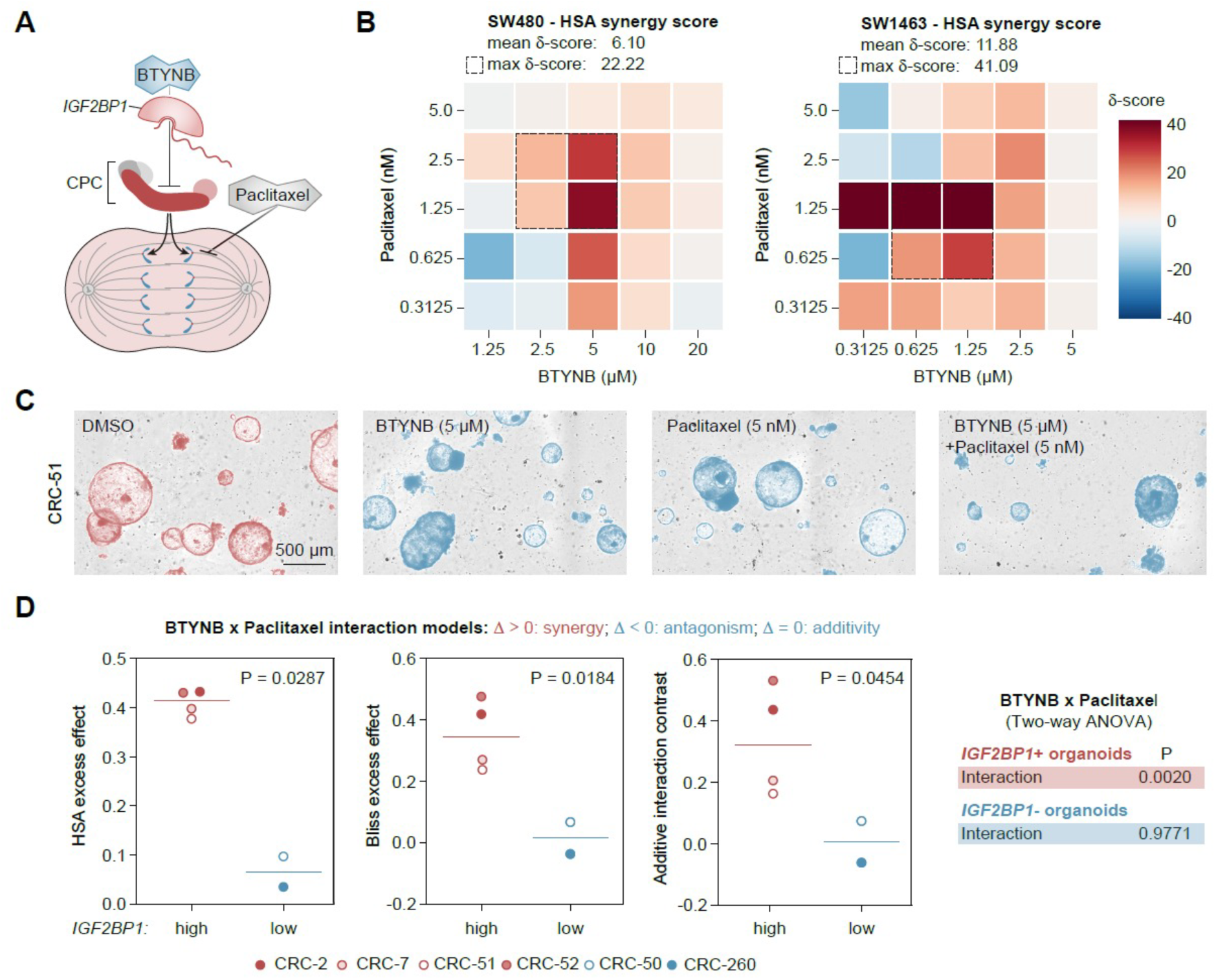
BTYNB synergizes with paclitaxel in CRC cells and IGF2BP1-positive organoids. **(A)** Schematic model illustrating how BTYNB-mediated inhibition of IGF2BP1 reduces chromosomal passenger complex (CPC) expression, while paclitaxel stabilizes microtubules, jointly disrupting mitotic spindle function. **(B)** BTYNB–paclitaxel interactions in SW480 (left) and SW1463 (right) cells treated for 48 h using 5 × 5 concentration matrices. Cell viability was measured by CellTiter-Glo (triplicate measurements per combination). Heatmaps show Highest Single Agent (HSA) δ-scores (red, synergy; blue, antagonism). Dashed boxes indicate the most synergistic 2 × 2 regions. Mean δ-score represents the average across the entire matrix; maximum δ-score indicates the highest individual score. **(C)** Representative images of CRC51 organoids following 72 h treatment with DMSO, BTYNB (5 µM), paclitaxel (5 nM), or their combination. Scale bar, 500 µm. **(D)** Comparison of BTYNB–paclitaxel combination effects between IGF2BP1-positive (CRC2, CRC7, CRC51, CRC52) and IGF2BP1-low (CRC50, CRC260) organoids using HSA excess effects (left), Bliss excess effects (middle), and inverted additive interaction contrasts derived from two-way ANOVA (right). Each point represents the mean of three independent experiments for one organoid line (n = 4 positive and n = 2 low); horizontal lines indicate group means. Positive values indicate effects exceeding the respective reference model. Group differences were assessed using unpaired two-tailed Welch’s t-tests. The accompanying table shows BTYNB × paclitaxel interaction P values from two-way ANOVA for IGF2BP1-positive and -low organoids. Individual treatment responses, pairwise comparisons using Šídák’s multiple comparisons test, and organoid-specific HSA and Bliss analyses are shown in Supplementary Fig. S4.

We next investigated whether this combination also enhances treatment responses in patient-derived CRC organoids. Using BTYNB concentrations approximately two- to threefold below the respective EC₅₀ values, we observed that BTYNB or paclitaxel alone moderately reduced organoid growth in IGF2BP1-positive CRC51 organoids, whereas combined treatment markedly decreased organoid number and size (Fig. 6C). Consistently, the combination reduced viability to approximately 30–45% of control across all four IGF2BP1-positive organoid lines (CRC2, CRC7, CRC51, and CRC52), exceeding the effects of either single agent (Supplementary Fig. S4A). Two-way ANOVA revealed significant BTYNB × paclitaxel interactions in all four lines and across the IGF2BP1-positive group (Fig. 6D; Supplementary Fig. S4A). Moreover, both HSA and Bliss analyses yielded positive synergy scores in all four IGF2BP1-positive organoid lines, although not every line–model comparison remained significant after Holm correction (Fig. 6D; Supplementary Fig. S4B).

In contrast, organoids with low or undetectable IGF2BP1 expression showed little additional benefit from BTYNB. In CRC50, paclitaxel reduced viability, but its effect was not further enhanced by BTYNB, whereas CRC260 showed only modest responses to either single agent or their combination (Supplementary Fig. S4A). Despite treatment with a higher BTYNB concentration, neither of the IGF2BP1-low lines showed a significant treatment interaction or evidence of synergy in the HSA and Bliss analyses (Fig. 6D; Supplementary Fig. S4). Importantly, the magnitude of the combination effect consistently distinguished IGF2BP1-positive from IGF2BP1-low organoids across all three interaction models (Fig. 6D). Together, these findings establish preferential BTYNB–paclitaxel synergy in IGF2BP1-expressing CRC organoids and highlight IGF2BP1 expression as a promising marker of combination treatment sensitivity.

## Discussion

This study identifies IGF2BP1 as a post-transcriptional regulator of the chromosomal passenger complex (CPC) and establishes a link between oncogenic RNA regulation and mitotic integrity. Genetic and pharmacological perturbation of IGF2BP1 impaired CPC expression, disrupted spindle organization and reduced cancer cell viability. Importantly, BTYNB-mediated inhibition sensitized CRC cells to paclitaxel, with preferential synergy in IGF2BP1-expressing patient-derived organoids. These findings reveal a therapeutically accessible mitotic vulnerability and extend the oncogenic functions of IGF2BP1 beyond its established regulation of proliferative signaling.

IGF2BP1 promotes cancer cell proliferation through stabilization of oncogenic transcripts, including MYC, E2F1 and YAP1 (13–15). Our findings uncover CPC regulation as an additional mechanism supporting successful cell division. Although BIRC5 has previously been identified as an IGF2BP1 target (35), the coordinated regulation of all four CPC components—INCENP, AURKB, CDCA8 and BIRC5—has not been established. Their recurrent downregulation following IGF2BP1 depletion across 15 cancer cell models representing different tumor entities, together with concordant protein changes in PANC-1 cells, suggests that this regulatory network extends beyond CRC. IGF2BP1 association with all four CPC transcripts was supported by eCLIP and validated by RIP in CRC cells. Notably, BTYNB impaired binding to INCENP, CDCA8 and BIRC5 and accelerated INCENP mRNA decay, providing mechanistic evidence for post-transcriptional control. INCENP is particularly relevant as the central CPC scaffold, and its depletion reproduced the pronounced spindle defects and loss of viability observed upon IGF2BP1 perturbation. Thus, IGF2BP1-dependent CPC regulation may represent a broadly conserved mechanism sustaining mitotic integrity in cancer cells.

The CPC coordinates chromosome segregation, spindle checkpoint signaling and cytokinesis (36–39), making its regulation particularly relevant to tumor cells undergoing rapid proliferation. The pronounced spindle abnormalities, G2/M accumulation and increased DNA content following BTYNB treatment reveal a mitotic consequence of IGF2BP1 inhibition that complements its established role in G1/S progression. While transient IGF2BP1 depletion predominantly affected G1/S distribution, genetic knockout and pharmacological inhibition produced more pronounced mitotic phenotypes. Importantly, spindle abnormalities were observed following all three perturbations, supporting a functional contribution of IGF2BP1 to mitotic organization. These findings suggest that IGF2BP1 supports not only proliferative entry through established MYC/E2F-associated programs but also successful mitotic progression through regulation of CPC expression.

The therapeutic relevance of this mechanism is supported by the treatment combination experiments. BTYNB and paclitaxel showed concentration-dependent HSA synergy in two CRC cell lines, consistent with increased vulnerability to microtubule-directed stress following disruption of CPC regulation. More importantly, all four IGF2BP1-positive patient-derived organoid lines displayed pronounced combination responses, whereas two IGF2BP1-low lines showed limited additional benefit from BTYNB despite treatment at higher concentrations. Concordant HSA, Bliss and ANOVA interaction analyses supported preferential combination effects in IGF2BP1-expressing organoids. These findings nominate IGF2BP1 expression as a candidate marker of combination sensitivity and provide a rationale for expression-guided targeting of mitotic vulnerabilities. Although paclitaxel is not an established CRC treatment, it serves as a proof-of-concept partner for exploiting this mechanism. Other mitosis-directed agents, including PLK1 and AURKB inhibitors, may offer opportunities for future therapeutic combinations (40–42).

Several limitations should be considered. Although the recurrent regulation of CPC transcripts across diverse cancer models suggests broader relevance, the functional consequences of this axis were investigated primarily in CRC, and its contribution to mitotic integrity in other tumor entities remains to be established. Furthermore, BTYNB has micromolar potency and incompletely defined target selectivity; additional compound activities may contribute to its effects, particularly given differences between acute pharmacological inhibition and genetic depletion. Finally, the organoid panel comprised four IGF2BP1-positive and two low-expressing models, limiting conclusions about the predictive value of IGF2BP1 expression. Larger, molecularly characterized organoid collections, genetic validation of treatment sensitivity and improved IGF2BP1-directed compounds will be important for advancing this therapeutic strategy.

## Conclusions

Our findings identify IGF2BP1-dependent CPC regulation as a previously unrecognized mechanism supporting mitotic integrity in cancer cells. Pharmacological disruption of this RNA-regulatory network induces spindle defects and sensitizes CRC cells to paclitaxel, with preferential synergy in IGF2BP1-expressing patient-derived organoids. Together, these results establish a mechanistic and translational rationale for exploiting IGF2BP1-associated mitotic vulnerabilities through expression-guided combination therapies.

## Declarations

### Ethics approval and consent to participate

Animals were handled according to the guidelines of the Martin Luther University. Permission was granted by the state administration office of Saxony-Anhalt (42502-2-1625 MLU).

### Consent for publication

Not applicable

### Availability of data and materials

The data created by RNA sequencing are deposited under GEO Accession: GSE350795 (https://www.ncbi.nlm.nih.gov/geo/query/acc.cgi?acc=GSE350795).

*will be released upon publication*

The mass spectrometry proteomics data have been deposited to the ProteomeXchange Consortium via the PRIDE [1] partner repository with the dataset identifier PXD085755.

*will be released upon publication*

### Authors’ contributions

BG, NB and SHü conceived and coordinated the project; TM, MH and LPM provided supervisory support; BG, NB and SHü contributed to data analysis and manuscript writing; BG, CM and AWG designed and performed cellular and in vitro experiments; NB and AR performed immunofluorescence analyses, cell cycle profiling, and organoid experiments and analyses; SHü, NB, DM and RW performed bioinformatics analyses; JL, TM and SHa designed and performed in vivo experiments; KUIM performed samples preparation for MS analyses, CI and FWO acquired mass spectrometric data acquisition and performed data analysis; CI and AS supervised MS analyses; SHü, JHK and NB contributed to funding acquisition. All authors read and approved the final manuscript.

## Acknowledgements

We thank the members of the DFG Research Unit FOR5433 for their valuable input and helpful discussions. The authors thank the Core Facility Imaging and the Core Unit NGS Analytics of the Medical Faculty, Martin Luther University Halle-Wittenberg, for providing equipment and technical support for imaging, flow cytometry and RNA-sequencing data processing.

## List of abbreviations

AURKB: Aurora Kinase B
BIRC5: Baculoviral IAP Repeat Containing 5 (Survivin)
BSA: Bovine serum albumin
BTYNB: Small-molecule inhibitor of IGF2BP1–RNA binding (2-[(5-CDCA8 bromothiophen-2-yl)methylideneamino]benzamide)
Cell: Division Cycle Associated 8 (Borealin)
COAD: Colon Adenocarcinoma
CPC: Chromosomal Passenger Complex
CRC: Colorectal carcinoma
DAPI: 4ʹ,6-diamidino-2-phenylindole
DMSO: Dimethyl sulfoxide
eCLIP: Enhanced crosslinking and immunoprecipitation
FBS: Fetal bovine serum
FDR: False discovery rate
FOLFIRI: Folinic acid, fluorouracil, irinotecan
FOLFOX: Folinic acid, fluorouracil, oxaliplatin
GFP: Green fluorescent protein
GO-BP: Gene Ontology Biological Process
GSEA: Gene set enrichment analysis
HSA: Highest Single Agent (synergy model)
IGF2BP1 / I1: Insulin-like growth factor 2 mRNA-binding protein 1
IF: Immunofluorescence
INCENP: Inner Centromere Protein
I1-KO: IGF2BP1 knockout
I1-OE: IGF2BP1 overexpression
I1-RE: IGF2BP1 re-expression
LFQ: Label-free quantification
MFI: Mean fluorescence intensity
MOI: Multiplicity of infection
MSigDB: Molecular Signatures Database
NES: Normalized enrichment score
PBS: Phosphate-buffered saline
PLK1: Polo-like kinase 1
RBP: RNA-binding protein
RIP: RNA co-immunoprecipitation
RT-qPCR: Reverse transcription quantitative PCR
SBP: Streptavidin-binding peptide
siC: Non-targeting control siRNA
si_I1: IGF2BP1-targeting siRNA
si_INCENP: INCENP-targeting siRNA
siRNA: Small interfering RNA
TCGA-COAD: The Cancer Genome Atlas — Colon Adenocarcinoma
TMM: Trimmed mean of M-values (normalization method)
TPM: Transcripts per million

