## Supplementary Figures for "Targeting IGF2BP1 Creates a Mitotic Vulnerability in Colorectal Carcinoma"

### Supplementary Material

Supplementary Figure S1

Supplementary Figure S2

Supplementary Figure S3

Supplementary Figure S4

### Supplementary Figure 1

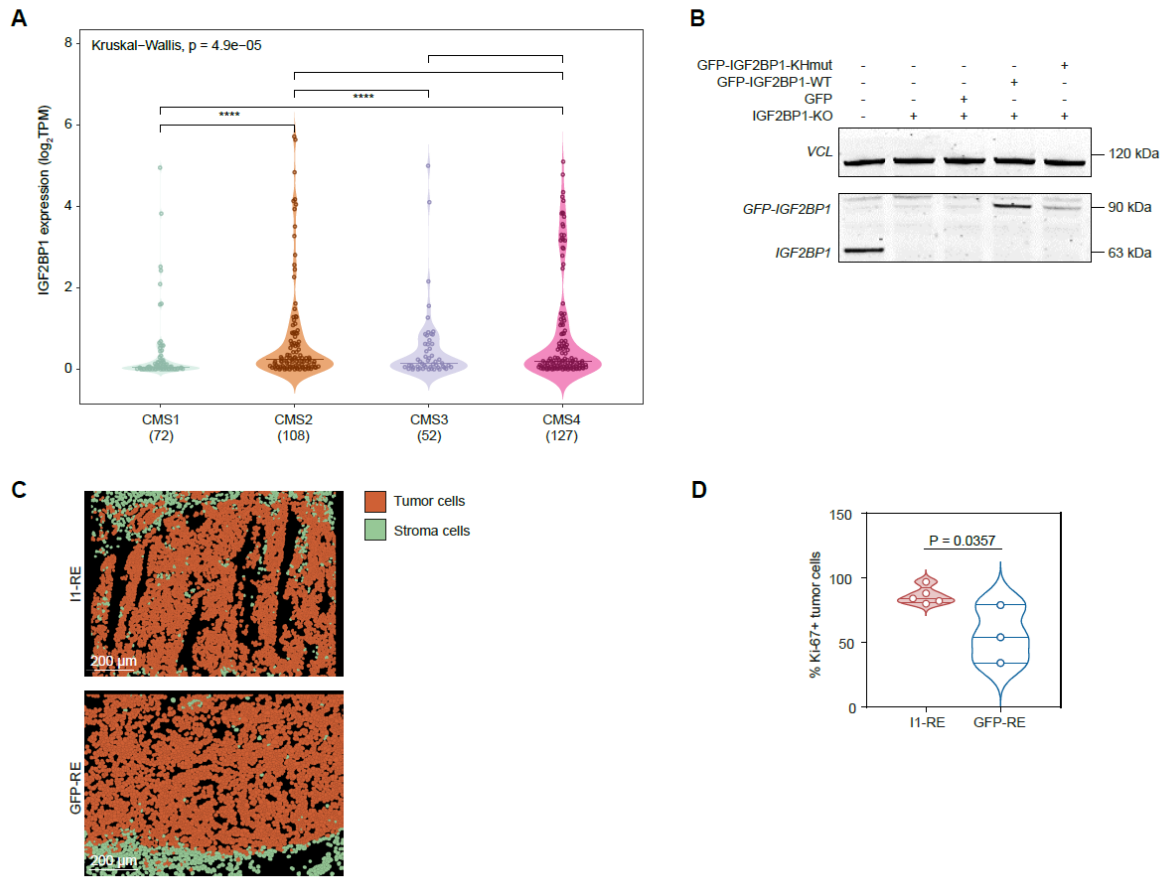

**Supplementary Fig. S1. IGF2BP1 expression across CRC molecular subtypes and characterization of xenograft models.** **(A)** IGF2BP1 mRNA expression across consensus molecular subtypes (CMS1–4) in the TCGA-COAD cohort ($n = 397$). Group differences were assessed using the Kruskal–Wallis test followed by multiple comparisons. **(B)** Western blot analysis of IGF2BP1 in SW1463 wild-type (WT), IGF2BP1-knockout (I1-KO), and I1-KO cells re-expressing GFP (GFP-RE), GFP-IGF2BP1 (I1-RE), or RNA-binding-deficient GFP-IGF2BP1-KHmut. Vinculin (VCL) serves as a loading control. **(C)** Representative segmentation masks distinguishing tumor cells (orange) from stromal cells (green) in xenografts derived from I1-RE and GFP-RE cells. Scale bars, 200 $\mu\text{m}$. **(D)** Percentage of Ki67-positive tumor cells in I1-RE and GFP-RE xenografts. Statistical significance was assessed using an unpaired two-tailed t-test. P values are indicated.

### Supplementary Figure 2

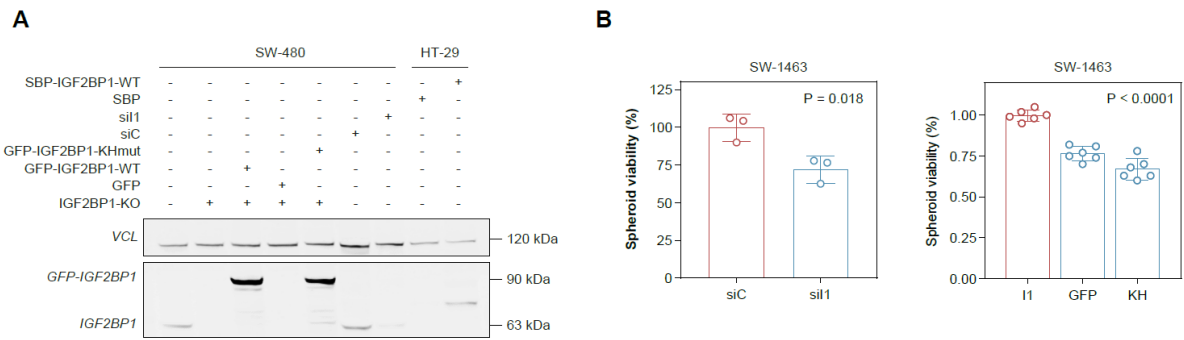

**Supplementary Fig. S2. Characterization of IGF2BP1-perturbed CRC cell models and their effects on spheroid viability. (A)** Western blot analysis of IGF2BP1 in SW480 wild-type, IGF2BP1-knockout (I1-KO), and I1-KO cells re-expressing GFP, GFP-IGF2BP1-WT, or RNA-binding-deficient GFP-IGF2BP1-KHmut, as well as SW480 cells following control (siC) or IGF2BP1-targeting (si1) siRNA transfection and HT29 cells with inducible IGF2BP1 expression. Vinculin (VCL) serves as a loading control. **(B)** Spheroid viability of SW1463 cells following IGF2BP1 knockdown (si1 versus siC; left) or re-expression of GFP-IGF2BP1 (I1), GFP alone, or GFP-IGF2BP1-KHmut in IGF2BP1-knockout cells (right). Data are shown as individual values with mean ± SD. P values are indicated.

**Supplementary Figure 3**

**A**

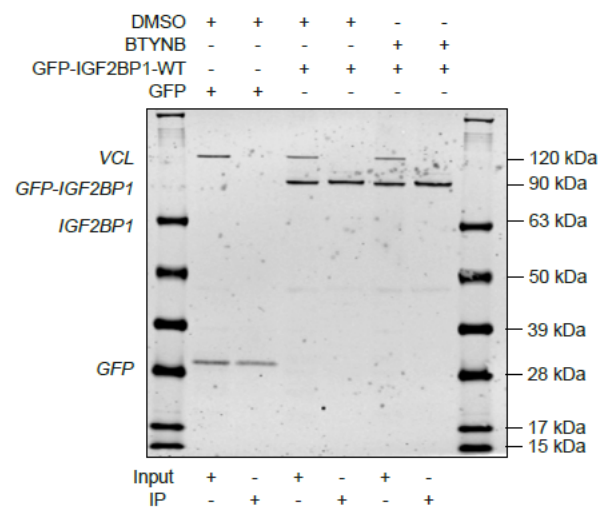

**Supplementary Fig. S3. Validation of GFP-IGF2BP1 immunoprecipitation following BTYNB treatment.** (A) Western blot analysis of input and GFP immunoprecipitation (IP) fractions from SW480 IGF2BP1-knockout cells re-expressing GFP or GFP-IGF2BP1, treated with DMSO or BTYNB (5 μM) for 24 h. GFP-IGF2BP1 and GFP were detected using an anti-GFP antibody. Vinculin (VCL) serves as an input control. Corresponding RNA immunoprecipitation results are shown in Fig. 5A.

### Supplementary Figure 4

**A**

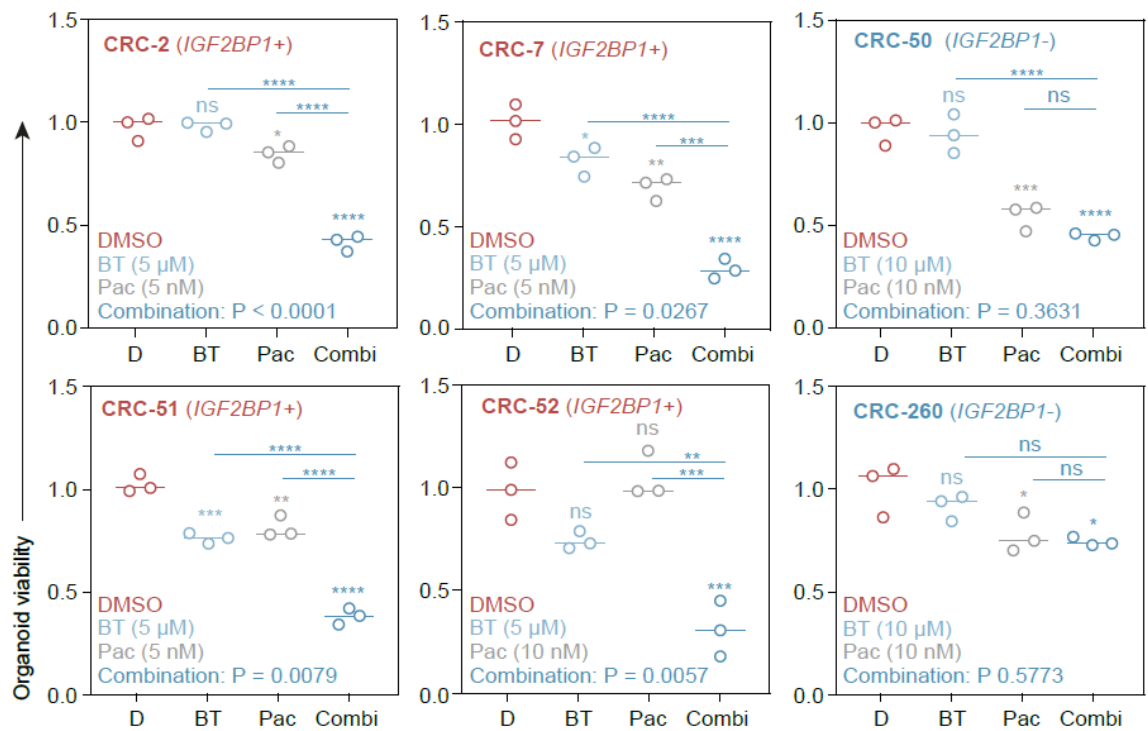

**B**

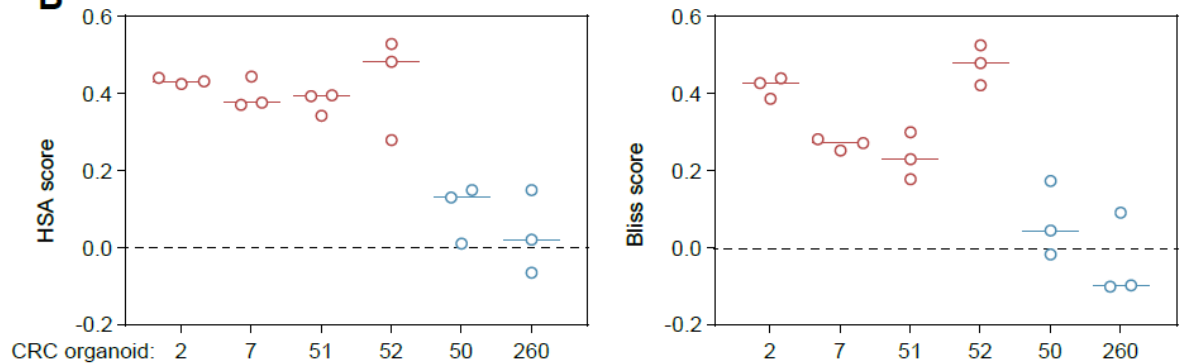

| Organoid | HSA |  | Bliss |  | IGF2BP1 expression |
| --- | --- | --- | --- | --- | --- |
|  | P-value | Holm-adj. P-value | P-value | Holm-adj. P-value |  |
| CRC2 | 0.0001 | 0.0006 | 0.0015 | 0.0075 | high |
| CRC7 | 0.0035 | 0.0140 | 0.0010 | 0.0060 | high |
| CRC51 | 0.0021 | 0.0105 | 0.0217 | 0.0651 | high |
| CRC52 | 0.0304 | 0.0912 | 0.0040 | 0.0160 | high |
| CRC50 | 0.1541 | 0.3082 | 0.3549 | 0.7098 | low |
| CRC260 | 0.6312 | 0.6312 | 0.6255 | 0.7098 | low |

**Supplementary Fig. S4. BTYNB and paclitaxel combination responses in patient-derived CRC organoids. (A)** Viability of four IGF2BP1-positive (CRC2, CRC7, CRC51, CRC52) and two IGF2BP1-low (CRC50, CRC260) organoid lines following treatment with DMSO, BTYNB (BT), paclitaxel (Pac), or their combination (Combi). Drug concentrations are indicated in each panel. Viability was normalized to DMSO controls. Statistical significance was assessed using two-way ANOVA with BTYNB × paclitaxel interaction terms and Šídák's multiple comparisons test. Interaction P values are indicated in each panel. **(B)** Highest Single Agent (HSA) and Bliss synergy scores calculated for the indicated drug combinations in each organoid line. Positive scores indicate effects exceeding the respective reference model; the dashed line indicates zero synergy. Individual points represent three independent experiments ($n = 3$); horizontal lines indicate medians. Statistical significance of HSA and Bliss scores was assessed using two-tailed one-sample t-tests against zero, with Holm correction for multiple testing across the six organoid lines within each model. Unadjusted and Holm-adjusted P values are provided in the accompanying table.
